# Feasibility evaluation of transtympanic laser stimulation of the cochlea from the outer ear

**DOI:** 10.1101/2021.11.08.467672

**Authors:** Miku Uenaka, Hidekazu Nagamura, Shizuko Hiryu, Kohta I. Kobayasi, Yuta Tamai

**Affiliations:** Graduate School of Life and Medical Sciences, Doshisha University, Kyotanabe, Kyoto, Japan; Organization for Research Initiatives and Development, Doshisha University, Kyotanabe, Kyoto, Japan

**Keywords:** infrared laser stimulation, auditory prosthesis, compound action potentials, tympanic membrane, optoacoustic effect

## Abstract

Infrared laser stimulation has been studied as an alternative approach to auditory prostheses. This study evaluated the feasibility of infrared laser stimulation of the cochlea from the outer ear bypassing the middle ear function. An optic fiber was inserted into the ear canal and a laser was used to irradiate the cochlea through the tympanic membrane in Mongolian gerbil. A pulsed infrared laser (10.1 mJ/cm^2^) and clicking sound (70 peak-to-peak equivalent sound pressure level) were presented to the animals. The amplitude of the laser-evoked cochlear response was systematically decreased following insertion of a filter between the tympanic membrane and cochlea; however, the auditory-evoked cochlear response did not decrease. The filter was removed and the laser-evoked response returned to around the original level. The amplitude ratio and the relative change in response amplitude before and during filter insertion significantly decreased as the absorbance of the infrared filter increased. These results indicate that laser irradiation could bypass the function of the middle ear and directly activate the cochlea. Therefore, an auditory prosthesis based on laser stimulation represents a possible noninvasive alternative to conventional auditory prostheses requiring surgical implants.

## 1. Introduction

More than 5% of the world’s population (around 466 million people) have communication difficulties caused by hearing loss (World Health Organization, 2018). Hearing aids are one of the most common auditory prostheses used to compensate for hearing problems; however, they require an intact middle ear and patients with middle ear abnormalities require an auditory prosthesis (e.g., bone-anchored hearing devices, middle ear implants, or cochlear implants) that bypass the function of the middle ear to stimulate the auditory system. One of the main obstacles to the widespread use of these devices is the requirement for implant surgery, which is associated with potential complications (Ahmed et al., 2021; Håkansson et al., 2008; Yeung et al., 2018).

Over the last decade, the application of infrared laser stimulation to auditory prostheses has been considered an alternative to current hearing devices (Littlefield and Richter, 2021). Laser stimulation activates spatially selective cell populations in a noncontact manner and does not require the delivery of exogenous agents to tissues, such as a viral vector; therefore, auditory prostheses with laser stimulation could replicate precise spectral information less invasively (Fekete et al., 2020). Although many studies have focused on the development of intracochlear stimulation systems to improve the spectral resolution of cochlear implants (Matic et al., 2013; Rajguru et al., 2010; Xu et al., 2018), no studies, as far as we know, directly investigate its potential for reducing the invasiveness of surgical implantation.

This study evaluated the feasibility of the use of laser stimulation of the cochlea from the outer ear to compensate for auditory deficits. Since the tissue component of the tympanic membrane does not tend to absorb infrared light (Heimann et al., 2021; Sordillo et al., 2014), laser irradiation projected from the outer ear could penetrate the tympanic membrane and reach the cochlea. Thus, laser irradiation from the outer ear could stimulate the cochlea directly through the tympanic membrane. Transtympanic membrane laser stimulation can overcome the limitations of a current implantable auditory prosthesis; therefore, this study examined whether laser stimulation could elicit a cochlear neural response.

## 2. Materials and methods

### 2.1 Animals

Nine experimentally naive adult Mongolian gerbils (6 females, 3 males; *Meriones unguiculatus*) were used in this study. All experimental procedures were approved by the Animal Experimental Committee of Doshisha University.

### 2.2 Animal surgery

Surgery and electrophysiological recordings were performed under anesthesia using an intramuscular combined injection of ketamine (47 mg/kg) and xylazine (9.3 mg/kg). Maintenance doses of ketamine (17.5 mg/kg) and xylazine (7.0 mg/kg) were injected every 30–50 min or if the animals showed signs of increasing arousal (e.g., voluntary whisker movement). The temperature of the gerbils was maintained using heating pads placed beneath the animals.

A metal head post was attached to the dorsal side of the gerbils’ skulls using dental cement (Shofu, Kyoto, Japan) to stabilize the head. The left pinna was removed to provide a clear view of the tympanic membrane and the left bulla was exposed by making an incision from the shoulder to the jaw. A wide range hole was made on the bulla around 3.35 mm anterocaudally and 2.30 mm ventrodorsally to allow the infrared filter and silver electrode to easily access the middle ear. The silver electrode (Nilaco, Tokyo, Japan; diameter: 0.13; impedance <20 kΩ) was inserted into the hole and hooked onto the bony rim of the cochlear round window for recording cochlear responses. The electrode was stabilized on the hole’s edge with acrylic glue and dental cement. A reference electrode was placed on the skin of the shoulder, which was wetted using saline.

### 2.3 Auditory stimuli

An acoustic click (pulse width, 100 μs) was used as an auditory stimulus and presented 100 times via an electrical dome tweeter (FT28D, Fostex, Tokyo, Japan). The repetition rate of the auditory stimuli was 100 Hz. The tweeter, which was driven by an amplifier (A-10, Pioneer, Tokyo, Japan), was placed in front of the gerbil’s nostrils approximately 10 cm from the ear opening. The stimulus signal was synthesized using a 192 kHz sampling rate and produced by a digital-to-analog converter (Octa-Capture, Roland, Shizuoka, Japan). The peak-to-peak equivalent sound pressure level (dB peSPL) of the stimulus was adjusted to 70 dB peSPL using a microphone (Type 1, ACO Pacific, Inc., Tokyo, Japan).

### 2.4 Laser stimuli

A pulsed infrared laser (wavelength, 1,871 nm; duration, 100 μs) was used as a laser stimulus. The repetition rate of the laser stimuli was 100 Hz. The laser was presented 100 times via an optical fiber (diameter, 100 μm; NA, 0.22) and inserted mediolaterally into the left outer ear canal using a micromanipulator (MM-3, Narishige, Tokyo, Japan) as described in earlier reports (Tamai et al., 2019; Tamai et al., 2020). The tip of stimulation fiber was set at around 0.7 mm in front of the tympanic membrane to ensure that the cochlea was irradiated transtympanically from the outer ear (Fig. 1A). A pulsed infrared laser was created using a diode laser stimulation system (BWF-OEM-1850, B&W TEK, Delaware, USA) with a wavelength comparable to those previously reported (Baumhoff et al., 2019; Izzo et al., 2006; Izzo et al., 2007). The radiant energy for each pulsed laser was measured using a digital power meter (PM100D, Thorlabs, Tokyo, Japan) with a thermal power sensor (S302C, Thorlabs, Tokyo, Japan), which calibrated to 10.1 mJ/cm^2^. The voltage commands for laser stimuli were generated using a digital-to-analog converter (Octa-Capture, Roland, Shizuoka, Japan).

**Fig 1.**
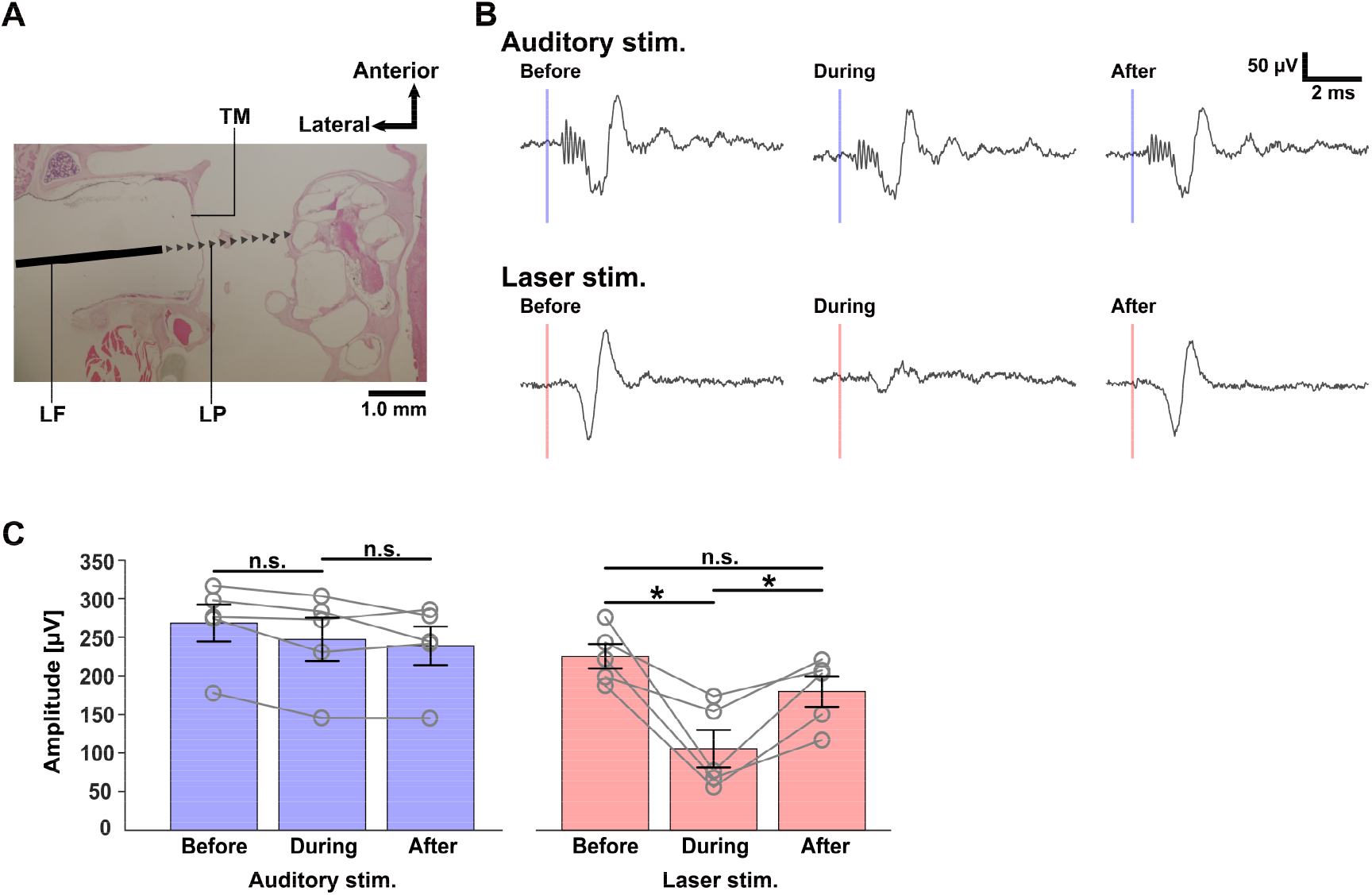
Effect of infrared filter insertion on auditory- and laser-evoked cochlear responses. (A) Horizontal section of a cochlea and reconstructed stimulation laser path. The locations of laser fiber (LF) and laser path (LP) were estimated based on our previous findings using microcomputed tomography images (Tamai et al., 2020), superimposed on hematoxylin and eosin-stained section. TM: tympanic membrane. (B) Cochlear response to auditory (100 μs single click with 70 dB peak-to-peak equivalent sound pressure level) and laser (100 μs single pulse, 10.1 mJ/cm^2^) stimuli before (left), during (center), and after (right) filter insertion (absorbance, 2.61). (C) Mean response amplitude evoked by auditory (blue) and laser (red) stimulation before, during, and after filter insertion. Open circles indicate the response amplitude in each animal. Error bars indicate the standard error of the mean (n = 5) * *P* < 0.05; two-tailed paired *t*-test with Bonferroni correction for multiple comparisons. n.s., not significant

### 2.5 Cochlear response recording before, during, and after filter insertion

An infrared filter with various absorbances (0.07, 2.61, and 3.22 at 1,871 nm) was inserted into the middle ear and held perpendicular to the laser optical path between the tympanic membrane and cochlea to modify the amount of laser irradiation reaching the cochlea without changing the radiant energy passing the tympanic membrane. The cochlear response elicited by auditory and laser stimulation was recorded before, during, and after filter insertion. The manipulation was intended to dissociate two possible stimulation sites (the tympanic membrane and cochlea) of the cochlear response evoked by transtympanic laser stimulation.

### 2.6 Measurement of thikness and optical absorbance of the tympanic membrane

The optical absorbance of the tympanic membrane was quantified in the gerbils. Four gerbils were euthanized and then decapitated and their tympanic bullae were extracted from the skulls. The tissues surrounding the tympanic membrane and auditory ossicles were carefully removed under a stereomicroscope. The thickness of the detached tympanic membrane was measured using a microcaliper (547-401, Mitutoyo, Kanagawa, Japan). Five points around the circumference of the midline between the umbo and annulus were measured at equal intervals within the ventral half of the pars tensa and the average thicknesses were calculated. The tympanic membrane was then placed on the glass slide and set on a thermal power sensor (S302C, Thorlabs, Tokyo, Japan) with a digital power meter (PM100D, Thorlabs, Tokyo, Japan) to measure the absorption. The laser fiber tip was directed vertically to the surface of the tympanic membrane and placed 0.7 mm from the tissue. The tympanic membrane on the glass slide was irradiated using the pulsed laser (duration, 100 μs; repetition rate, 4,000 Hz) at 10.1 mJ/cm^2^. The original radiant energy (I_0_) and transmitted energy of the tympanic membrane (I) was recorded to calculate the absorbance (A) as previously described (Goblet et al., 2021; Sorg et al., 2021) using the following equation:

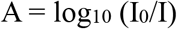

### 2.7 Data analyses

The cochlear microphonic (CM) and compound action potential (CAP) were separated according to their latency as described in earlier reports (Izzo et al., 2006; Tamai et al., 2019; Tamai et al., 2020; Teudt et al., 2011). The delays from stimulus onset to the first peak of the CM and CAP responses were measured. The amplitude was measured as the peak-to-peak voltage of each response and the amplitude ratio was evaluated as the relative response amplitude change before and during filter insertion. A two-tailed paired *t*-test with Bonferroni correction for multiple comparisons was conducted to assess the effect of filter insertion. Analyses were performed using either MATLAB (MathWorks, Natick, MA, USA) or R (R Core Team, 2018).

## 3. Results

Cochlear responses elicited by auditory (70 dB peSPL) and laser (10.1 mJ/cm^2^) stimulation are depicted in Fig. 1B. Prior to insertion of the filter, the auditory stimulus induced a CM of 0.49 ± 0.03 ms and CAP of 1.81 ± 0.08 ms (mean ± SEM, n = 5) after the onset of the stimulus, respectively, whereas laser stimulation evoked a CAP of 1.55 ± 0.07 ms after stimulus onset and did not induce a visually observable CM. The effect of the infrared filter (absorbance, 2.61) on the average CAP amplitude is shown in Fig. 1C. In the auditory-evoked response, the averaged CAP amplitude measurements before, during, and after filter insertion were 268.4 ± 24.0, 247.1 ± 28.0, and 238.9 ± 25.0 μV, respectively. In the laser-evoked response, the CAP amplitude decreased from 225.4 ± 15.9 to 105.6 ± 24.2 μV after insertion of the filter. The CAP amplitude returned to 179.6 ± 19.8 μV after the filter was removed. The laser-evoked response amplitude was significantly smaller during the procedure than before and after the procedure (*P* < 0.05, n = 5), whereas the auditory-evoked response amplitude was not significantly different before, during, or after the procedure (*P* > 0.12 for all). The CAP amplitude ratio at different filter absorbances before and during insertion of the filter (Fig. 2A) indicated that the amplitude ratio decreased significantly as the absorbance increased (r = −0.67, *P* < 0.05). Fig. 2B shows the correlation between the thickness of the tympanic membrane and the optical absorbance of infrared laser radiation (n = 7). The mean thickness was 18 ± 6 μm and the absorbance was 0.054 ± 0.01. The absorbance was significantly positively correlated with the thickness of the tympanic membrane (r = 0.76, *P* < 0.05).

**Fig 2.**
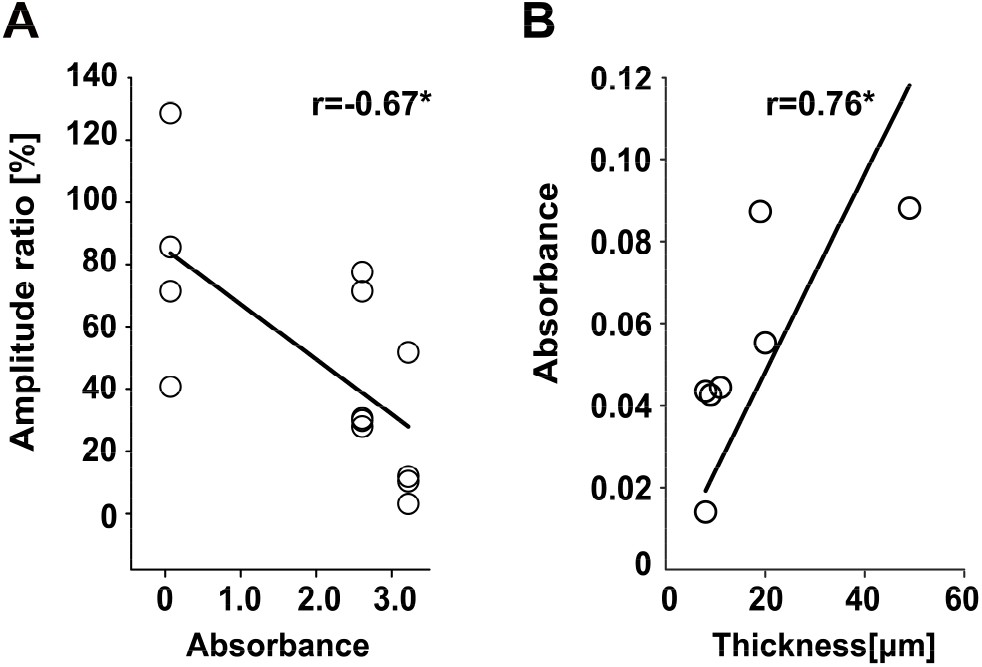
Change in amplitude ratio following filter insertion with different absorbance (A) and correlation between the thickness of the tympanic membrane and its absorbance (B). Data were obtained from 5 (A) and 7 (B) animals and correlation coefficients were calculated using Pearson’s correlation analysis. Open circles indicate individual data. * *P* < 0.05

## 4. Discussion

Our results showed that laser irradiation of the cochlea from the outer ear was able to elicit a CAP response similar to that of an auditory-evoked response (Fig. 1B). Teudt et al. (2011) demonstrated that infrared laser treatment using 50 mJ/cm^2^ irradiation evoked a sound of approximately 40 dB peSPL 2 mm from the glass fiber tip in the air, which suggests that care must be taken to distinguish between an optoacoustically induced response and an optoneural response. Therefore, it is possible that the laser vibrated the tympanic membrane either directly or thermoacoustically and produced a CAP. However, our findings suggest otherwise. We reduced the radiant energy reaching the cochlea without changing the energy passing through the tympanic membrane by inserting an infrared filter between the tympanic membrane and the cochlea. Our results showed that the laser-induced response was decreased by the infrared filter insertion whereas the auditory-induced response was not reduced (Fig. 1B and C). Moreover, the relative amplitude ratio was significantly negatively correlated with the absorbance of the filter (Fig. 2A). Taken together, the amplitude of the intensity of the cochlear response is dependent on the radiant energy reaching the cochlea but not the tympanic membrane. These findings suggest that laser stimulation from the outer ear can be used to stimulate the cochlea while bypassing the middle ear function.

It is important to ensure that the laser irradiation of the cochlea does not damage the tympanic membrane during transtympnaic stimulation. Several studies have directly assessed the biocompatibility of laser irradiation with the tympanic membrane in vivo (Sorg et al., 2019) and in vitro (Pillong et al., 2020). These studies suggested that thermal damage depends on optical absorption during the interaction between the laser and tissue and caused cell necrosis. In this study, the thickness of the tympanic membranes in the gerbils was 18 ± 6 μm, which is in line with the findings of previous studies (von Unge et al., 1991). Thicker tympanic membranes showed a significantly higher absorbance (Fig. 2B). Higher absorbance of the tympanic membrane requires more radiant energy to stimulate the cochlea. Since humans have a thicker tympanic membrane than gerbils (30– 90 μm; Lim, 1970), this could be associated with a greater risk of damage to the tympanic membrane. Although our previous study demonstrated that continuous transtympanic membrane laser irradiation at 12.8 mJ/cm^2^ for 1 h did not result in observable damage to the tympanic membrane in gerbils (Tamai et al., 2019), safety evaluation using a human tympanic membrane must be performed prior to the practical application of infrared laser stimulation in auditory prostheses.

Our data demonstrated that laser stimulation from the outer ear elicited auditory neural activities of the cochlea; however, the level of hearing impairment, for which this technique will be effective, remains unclear. Some studies have reported that laser-induced auditory responses were not observed after chemical hearing impairment (i.e., neomycin administration into the cochlea), which suggests that laser stimulation may only apply to individuals with mild-to-moderate hearing loss whose hair cells work at least partially (Baumhoff et al., 2019; Schultz et al., 2012; Thompson et al., 2015). However, Richter et al. (2008) reported that the laser-induced cochlear response did not significantly decrease after 30–40 dB chemical deafening of the auditory threshold. Tan et al. (2018) revealed that laser stimulation could elicit auditory neural activity in congenitally deaf mice in which the cochlea lacks synaptic transmission between the inner hair cells and spiral ganglion neurons. Additionally, the laser-induced response cannot be masked by auditory stimuli when animals have an impaired hearing threshold (Young et al., 2015) These results indicate that laser irradiation of the cochlea directly activates auditory nerves and bypasses hair cells. This notion is partially compatible with our findings (Fig. 1B) that show that cochlear laser stimulation can produce a CAP without a clear CM response, as reported in earlier studies (Izzo et al., 2006; Tamai et al., 2019; Tamai et al., 2020; Teudt et al., 2011). These findings suggest that laser treatment can produce an auditory response in individuals with profound hearing loss who require cochlear implants. Future studies are required to directly compare the effects of laser stimulation among individuals with different levels of impairment in the auditory periphery to evaluate the efficacy of this method.

## 5. Conclusion

This study examined the feasibility of laser irradiation from the outer ear. Our results showed that laser stimulation directly activated the cochlea, bypassed the function of the middle ear, and induced a neural response similar to an auditory-evoked response. These findings suggest that laser irradiation from the outer ear could compensate for conductive hearing loss. If the laser irradiation of the cochlea can directly activate cochlear nerves, as some research groups described (Richter et al., 2008; Tan et al., 2018; Young et al., 2015), the auditory prosthesis based on the laser stimulation from the outer ear may be a possible non-invasive alternative to electrical cochlear implants. Auditory prosthesis with infrared laser stimulation does not require surgery and could help many hard-of-hearing individuals to sense sounds.

## Acknowledgments

We wish to thank Hiroshi Riquimaroux, Suguru Matsui, Tomohiro Miyasaka, and Kazuki Tanaka for their valuable advice regarding the experimental design. This research was supported by the Japan Society for the Promotion of Science (JSPS) KAKENHI Grants Nos. 21H03469 (K.I.K.), 21K21322 (Y.T.).

